# An end-to-end platform for pose estimation and real-time edge-AI deployment

**DOI:** 10.64898/2026.01.24.700912

**Authors:** David L. Haggerty, Caleb B. Darden, David M. Lovinger

## Abstract

Accurate pose estimation underpins quantitative analysis of behavior, yet many deep learning-based tracking tools remain optimized for offline workflows that rely on fragmented software pipelines, workstation-grade GPUs, or external middleware to enable real-time deployment. Here we present an integrated software-hardware ecosystem for pose estimation that spans dataset creation, model training, offline analysis, and real-time deployment on embedded edge-computing devices. SqueakPose Studio provides a software suite for whole-frame, deep learning-based pose estimation that unifies dataset creation, manual and model-assisted labeling, model training, validation, and large-scale offline inference. The system leverages modern object-detection architectures to enable efficient end-to-end training and inference without patch-based sampling or multi-stage postprocessing, and supports execution on CPUs, GPUs, and Apple Silicon. For experimental settings requiring continuous recording and synchronized data acquisition, SqueakView enables real-time model deployment, video capture, and sensor logging on embedded edge-computing hardware, while MouseHouse provides a compact, modular enclosure designed for home cage-based experiments that integrates embedded GPU compute, microcontroller-based timing, and peripheral I/O. A shared data format and deterministic timing architecture ensure consistency across offline analysis and real-time deployment. Together, SqueakPose Studio, SqueakView, and MouseHouse provide a unified platform for pose estimation that supports both conventional offline analysis and embedded, real-time experimentation, without reliance on workstation-grade hardware or external middleware.

## Introduction

Quantitative behavioral analysis is fundamental to neuroscience^1–12^, yet its implementation remains limited by the technical barriers of video annotation, computational throughput, and deployment scalability. In recent years, advances in deep learning have transformed pose estimation, enabling precise tracking of animal movements without physical markers. Software tools such as DeepLabCut,^13^ SLEAP,^14^ and Lighting Pose^15^ have established powerful foundations for markerless landmark tracking using convolutional neural networks trained on small, manually labeled datasets. These systems – initially built on ResNet-based^16^ architectures and more recently incorporating lighter and transformer-style backbones – achieved human-level accuracy in pose tracking through transfer learning with modest data volumes, particularly in rodent behavioral research. However, their reliance on patch-based sampling, multi-stage postprocessing, and external middleware such as Bonsai^17^ for real-time^18^ deployment introduces variable latency and complicates deterministic synchronization across devices, limiting their suitability for embedded or high-throughput experiments.

As behavioral paradigms grow in scale and complexity, there is an increasing need for analysis tools that are not only accurate but also fast, modular, and deployable on affordable edge hardware. Traditional pose-estimation pipelines were designed primarily for offline analysis, making them poorly suited for continuous, multimodal experiments that integrate video, sensor, and behavioral data in real-time. Meeting this challenge requires models capable of whole-frame inference, minimal postprocessing, and direct interfaces for experimental control – features exemplified by modern object-detection architectures such as You Only Look Once (YOLO)^19–21^. YOLO’s transformer-augmented backbone, single-pass detection, and unified representation of bounding boxes, keypoints, and segmentation masks provide a robust foundation for real-time behavioral tracking without crop-based preprocessing.

To address these limitations, we developed SqueakPose Studio, a modular software-hardware ecosystem for high-speed behavioral pose estimation and analysis built on modern YOLO architectures. SqueakPose unifies data labeling, model training, and analysis within an integrated framework that supports multimodal annotation, CUDA GPU and Apple Silicon acceleration, and semi-automated behavior clustering via UMAP and HDBSCAN. SqueakPose Studio, streamlines dataset creation through assisted labeling and confidence-based frame sampling, while the SqueakPose Analysis Utility enables quantitative behavior mapping. Complementing this software stack, the MouseHouse hardware platform and SqueakView software suite combines real-time video acquisition with sensor logging through a Jetson Orin Nano Super and RP2040 microcontroller. This configuration enables millisecond-precision recording and on-device YOLO inference via TensorRT, supporting closed-loop control in which behavioral detections can update task contingencies or external hardware in real-time. An expandable GPIO architecture ensures precise synchronization across devices, facilitating integration with complementary neuroscience systems such as *in vivo* photometry or electrophysiology. Together, SqueakPose Studio, MouseHouse, and SqueakView provide an end-to-end, low-cost platform for scalable, real-time, embedded behavioral analysis in freely behaving animals.

## Results

SqueakPose Studio provides a unified, modular environment for behavioral estimation that links dataset creation, training, and inference within a single framework (Fig.1a). Raw videos, image sequences, or pre-existing annotations can be converted into YOLO-compatible datasets through standardized input–output modules. The integrated labeling interface, SqueakPose Studio, supports bounding-box and keypoint annotation, allowing manual or model-assisted labeling and automated generation of training and validation splits. These components feed directly into YOLO-based training modules for detection and pose estimation, each compatible with CPU and CUDA GPU or Apple MPS acceleration. The underlying YOLO architecture combines convolutional and transformer elements within a C2f-Darknet backbone and PAN-FPN++ feature-fusion neck, with dedicated heads for detection, pose, and segmentation tasks (Fig.1b). This end-to-end configuration eliminates the patch-cropping and multi-stage refinement steps required by earlier two-stage pose-estimation frameworks, streamlining both training and inference. When benchmarked against DeepLabCut (ResNet-50 backbone) under identical conditions and training sets, YOLOv11s-pose trained 4.5 times faster, representing a 77.9% reduction in total training time (Fig.1c). During inference on a ten-minute, top-down open-field video SqueakPose Studio preformed offline, full video prediction 8.5 times faster than DeepLabCut, corresponding to effective inference frame rates of 35 FPS and 4 FPS, respectively (Fig.1d). Despite the substantial gains in computational efficiency, both approaches produced similar spatial precision across six anatomical keypoints – nose, head, left ear, right ear, back, and tailbase – with largely overlapping pixel-error distributions (Fig.1e). For training loss and mean average precision (mAP) scores of both models, see Figure1-figure supplement 1. Together, these benchmarks demonstrate that SqueakPose Studio achieves DeepLabCut-level accuracy while providing an order-of-magnitude improvement in training and offline inference speed.

In addition to speed improvements, we sought to implement a graphical interface for dataset inspection, labeling, and model-assisted annotation that natively supports all modern operating systems. Built in Python 3.12 using the PyQt6 framework, Squeakpose Studio enables users to browse datasets, save annotations, generate predictions, and train models through a single control panel (Fig.2a). Multiple labeling modes support flexible annotation workflows (Fig.2b).

Bounding boxes can be drawn or resized with crosshair assistance, while individual keypoints can be placed and assigned visibility states (visible, occluded, or out-of-frame). Labels can be manually edited or refined using local model inference, which predicts keypoints on selected frames to accelerate dataset generation (Fig.2c). Dataset validation and export tools are integrated directly within the GUI (Fig. 2d). Users can automatically construct YOLO-compatible directory structures, generate .yaml configuration files, and define train/validation split ratios without command-line interaction. The model-loader panel allows pre-trained YOLO weights (.pt files) to be loaded for assisted labeling, new frame prediction, and final full video inference. A dedicated training interface provides configuration options for dataset selection, model size, automatic GPU detection, epoch count, and batch size (automatic on CUDA GPUs) (Fig.2f). For further information, including a comprehensive live video demonstration of all features, please consult the project’s GitHub repository. Together, these features make SqueakPose Studio a native cross-platform tool for managing the full lifecycle of behavioral pose-estimation datasets from annotation and validation to training and inference through a single, user-friendly interface.

**Figure 1 -.**
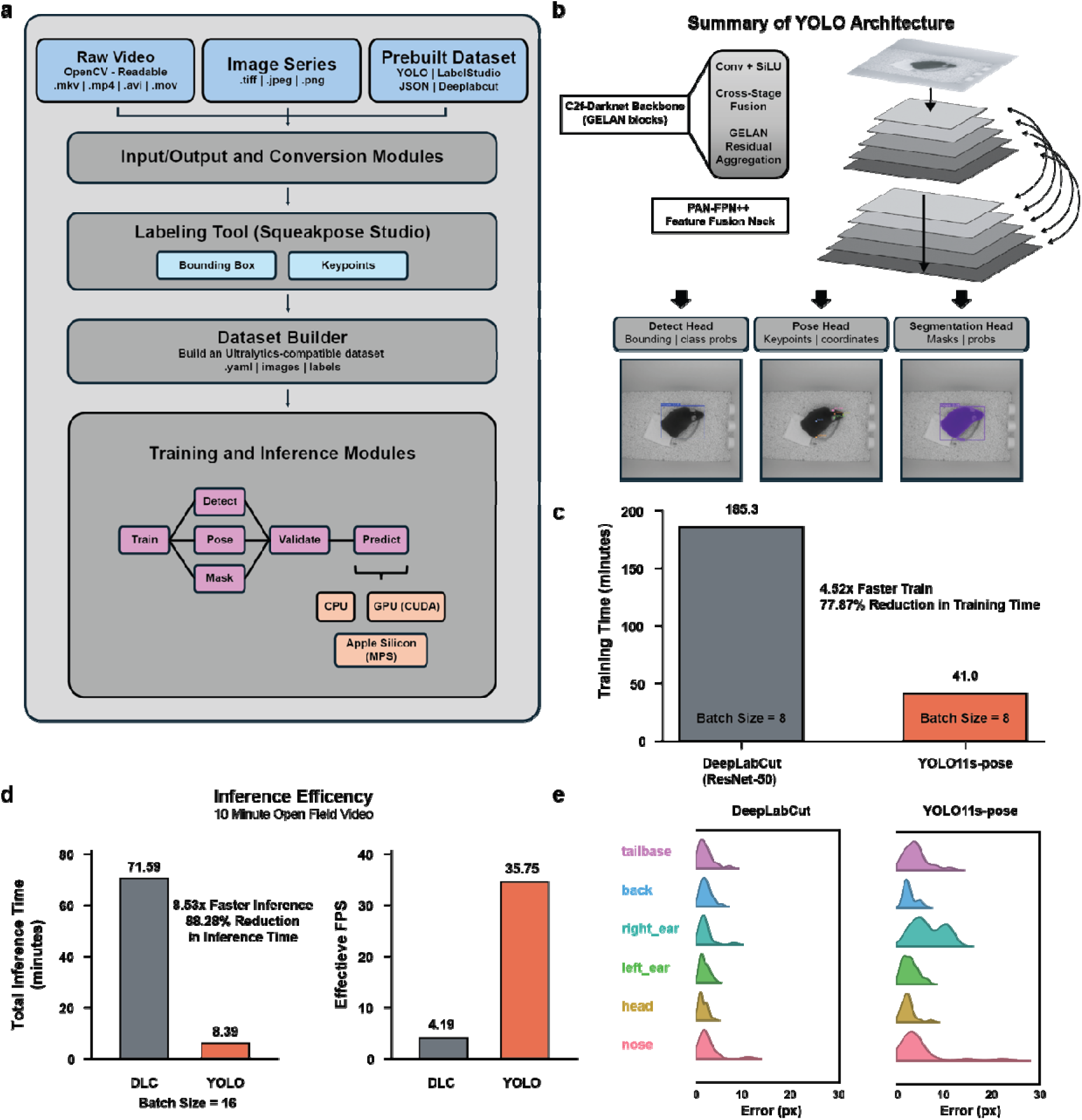
SqueakPose Studio provides a unified, modular environment for behavioral pose estimation that links dataset creation, training, and inference. (a) Software architecture linking video import, labeling, training, and analysis. (b) YOLOv11s-pose network design with C2f-Darknet backbone, PAN-FPN++ neck, and multitask detection, pose, and segmentation heads. (c) Training speed comparison between YOLOv11s-pose and DeepLabCut (ResNet-50) under identical dataset conditions. (d) Inference speed comparison on a 10-min open-field video showing an 8.5× improvement in processing rate. (e) Distribution of keypoint localization error across six anatomical landmark showing comparable spatial precision. Together, these benchmarks demonstrate that SqueakPose Studio achieves DeepLabCut level accuracy with markedly improved training and inference efficiency.

**Figure 2 -.**
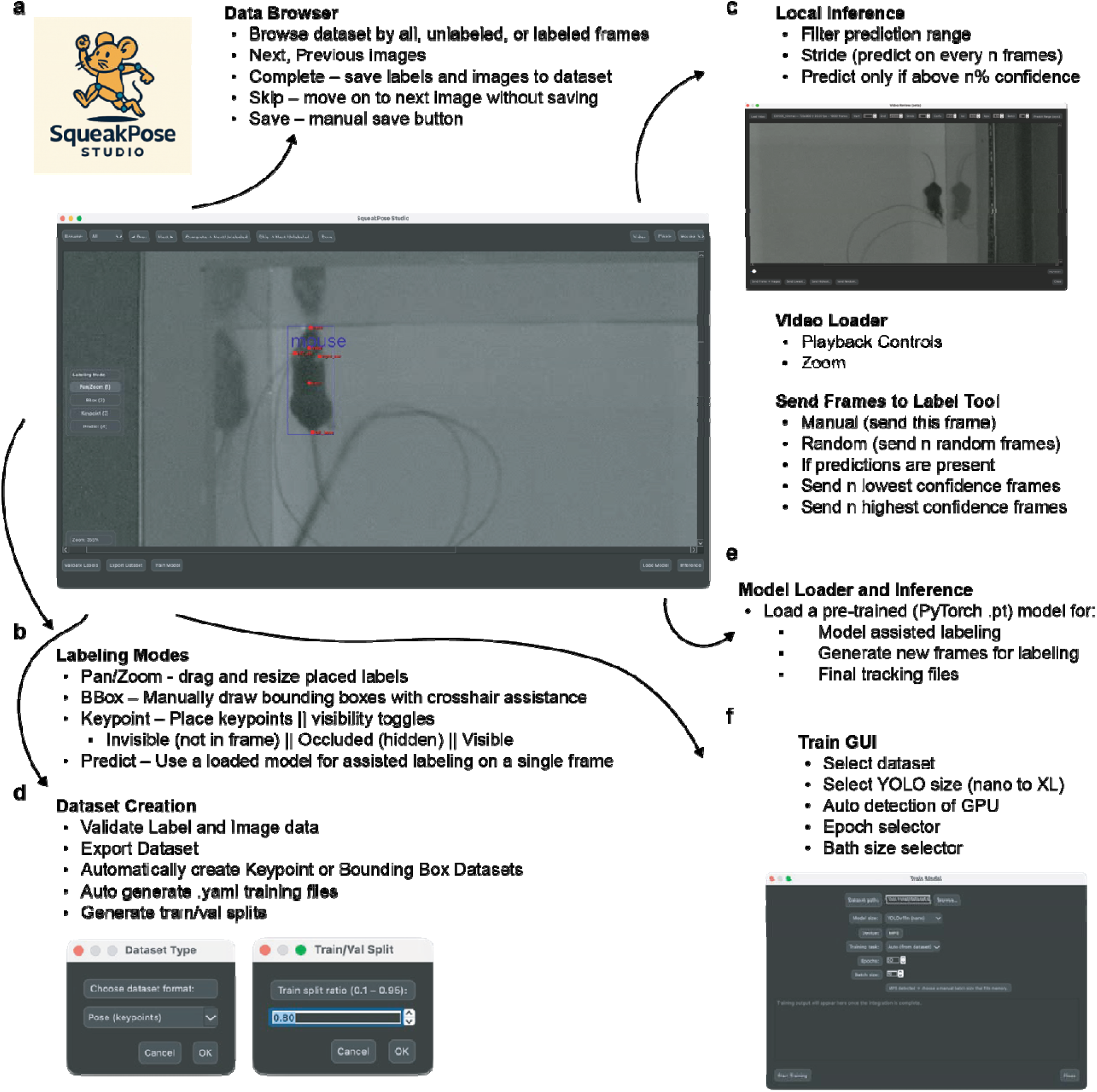
The SqueakPose Studio GUI supports dataset management, annotation, and model training across operating systems. **(a)** Application overview showing integrated dataset browser, model configuration, and training panels. **(b)** Bounding-box and keypoint labeling modes with crosshair placement and visibility-state assignment. **(c)** Model-assisted labeling using on-device YOLO predictions for annotation acceleration. **(d)** Dataset validation and export tools for generating YOLO-compatible structures and .yaml configuration files. **(e)** Integrated training interface for selecting model size, GPU, epochs, and batch size. Built in Python 3.12 with PyQt6, SqueakPose Studio enables interactive annotation and training without command-line dependencies.

**Figure 3 -.**
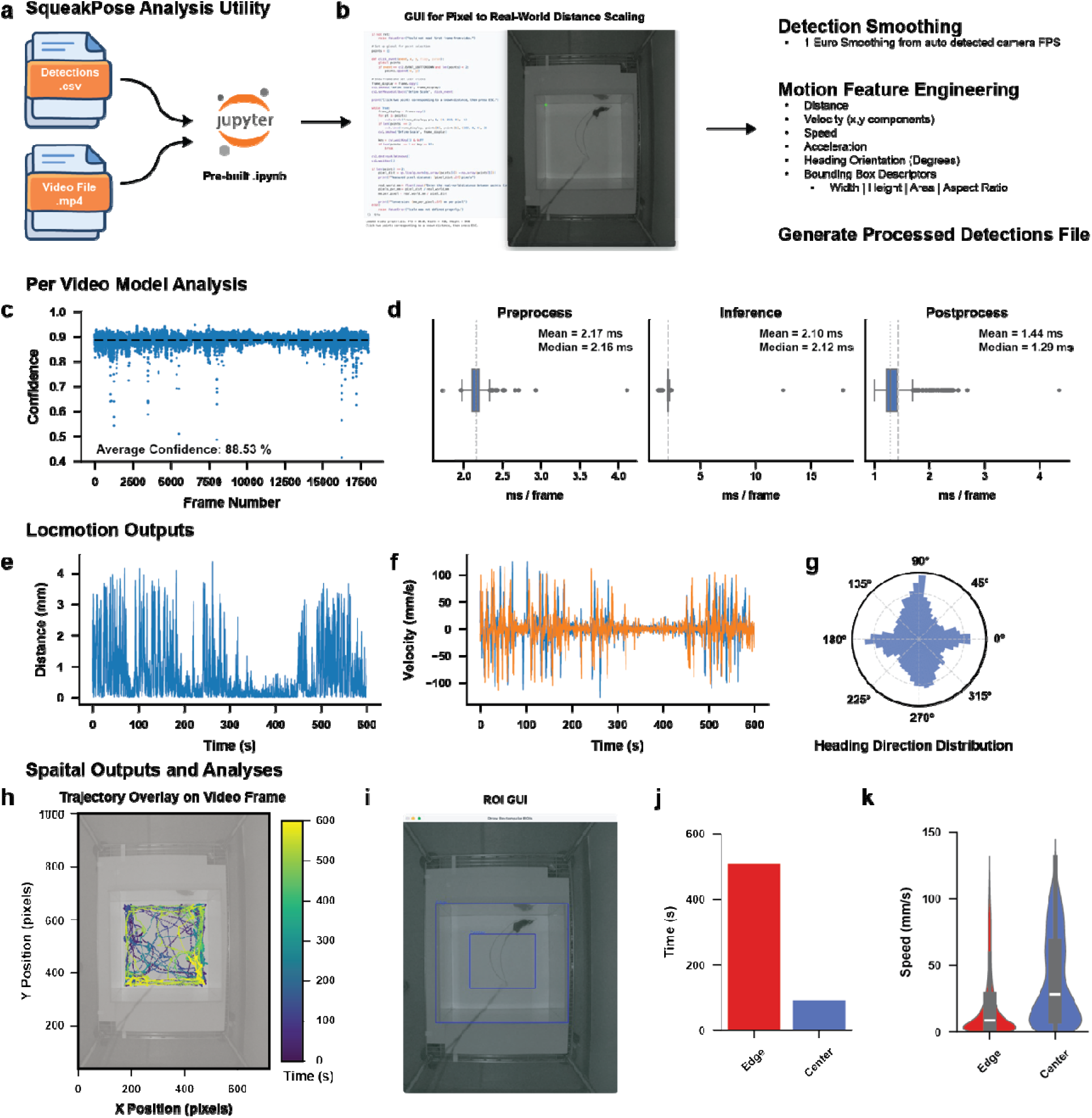
SqueakPose Analysis: A pre-built Jupyter notebook automates motion feature extraction and visualization from SqueakPose Studio detection outputs. **(a)** Workflow for loading detection (.csv) and vide (.mp4) files into the analysis utility. **(b)** Graphical interface for converting pixel coordinates to real-world units and applying 1-Euro smoothing; derived features include distance, velocity, acceleration, heading, and bounding-box descriptors. **(c)** Example model confidence trace across frames (mean ≈ 88.5%). **(d)** Frame-processing latency before, during, and after inference. **(e–f)** Locomotion profiles showing frame-wise distance and velocity over a 10-min open-field session. **(g)** Heading-direction distribution illustrating orientation sampling. **(h–i)** Spatial trajectory overlay and region-of-interest (ROI) GUI for defining arena zones. **(j–k)** Quantification of time and speed by ROI, revealing edge-biased exploration and higher center-zone velocities consistent with anxiogenic center behavior.

While existing tools can generate reliable tracking datasets from pose estimation, a substantial amount of time is often spent determining how to analyze these outputs. Recent utilities have been introduced to assist behavioral analysis from DeepLabCut and SLEAP tracking data^22–26^, yet these implementations typically require additional customization or scripting. To simplify this process, we developed an interactive python notebook that performs automated post-inference analysis of SqueakPose detections, providing a unified workflow for evaluating both model performance and basic behavioral metrics (Fig.3a). The notebook calculates confidence and timing statistics for each frame to assess inference stability and processing speed, and it automatically converts pixel-based detections into real-world coordinates using a built-in calibration GUI leveraging metadata from associated video files (Fig.3b). From these processed outputs, the utility derives standard motion features – including distance, velocity, acceleration, and heading orientation – along with spatial and region-of-interest analyses that quantify locomotion and exploration patterns (Fig.3c–k). The region of interest tool allows experimenters to simple annotate frames to create ROI masks that are then applied to the full dataset. In the example dataset shown here, a single animal was recorded in a top-down open-field assay, and SqueakPose Analysis recovered expected behavioral signatures commonly observed in open-field assays. The animal spent the greatest time along the perimeter of the assay (Fig.3j) and moved at higher speeds when traversing the center (Fig3.k), consistent with the interpretation that open-field center regions produce anxiogenic behaviors^27^. These results illustrate how the analysis utility can rapidly transform raw pose outputs into interpretable measures of locomotion and spatial preference, validating both model accuracy and behavioral relevance within a single, automated workflow.

The rapid adoption of computer vision–based pose estimation has enabled increasingly rich, high-dimensional datasets for behavioral neuroscience. To demonstrate how such data can be explored within the SqueakPose Studio framework, we implemented an example semi-supervised clustering workflow that links motion features to representative behavioral video segments (Fig.4a). This module extracts kinematic and spatial descriptors – including distance, velocity, acceleration, heading orientation, and temporal bins – from processed detections and embeds them into a low-dimensional space using UMAP (Fig.4b). Clusters are identified in this embedding using HDBSCAN, which automatically determines cluster membership based on local density, generating a set of candidate behavioral groups. To visualize relationships among these groups, the pipeline computes hierarchical cluster similarity using Ward’s linkage method (Fig.4c), enabling users to explore how behaviors segregate or merge across distance thresholds. For demonstration, we applied this workflow to the same open field recording and recovered several distinct pose-defined behavioral motifs, including edge exploration, edge stationary, and center exploration (Fig. 4d). A module in the analysis tool allows for recovering video clips belonging to identified clusters to allow researchers to define these groups in a supervised fashion. This allows parameter tuning for UMAP and HDBSCAN to occur in a closed-loop fashion to uncover human-understandable features. This module is designed as a flexible starting point for exploratory analysis and can be used to generate labeled datasets for downstream modeling. All processed outputs – including feature matrices, cluster assignments, and embeddings – can be exported for integration with other popular analytical tools such as CEBRA^28^ or Keypoint-MoSeq^29^.

**Figure. 4 -.**
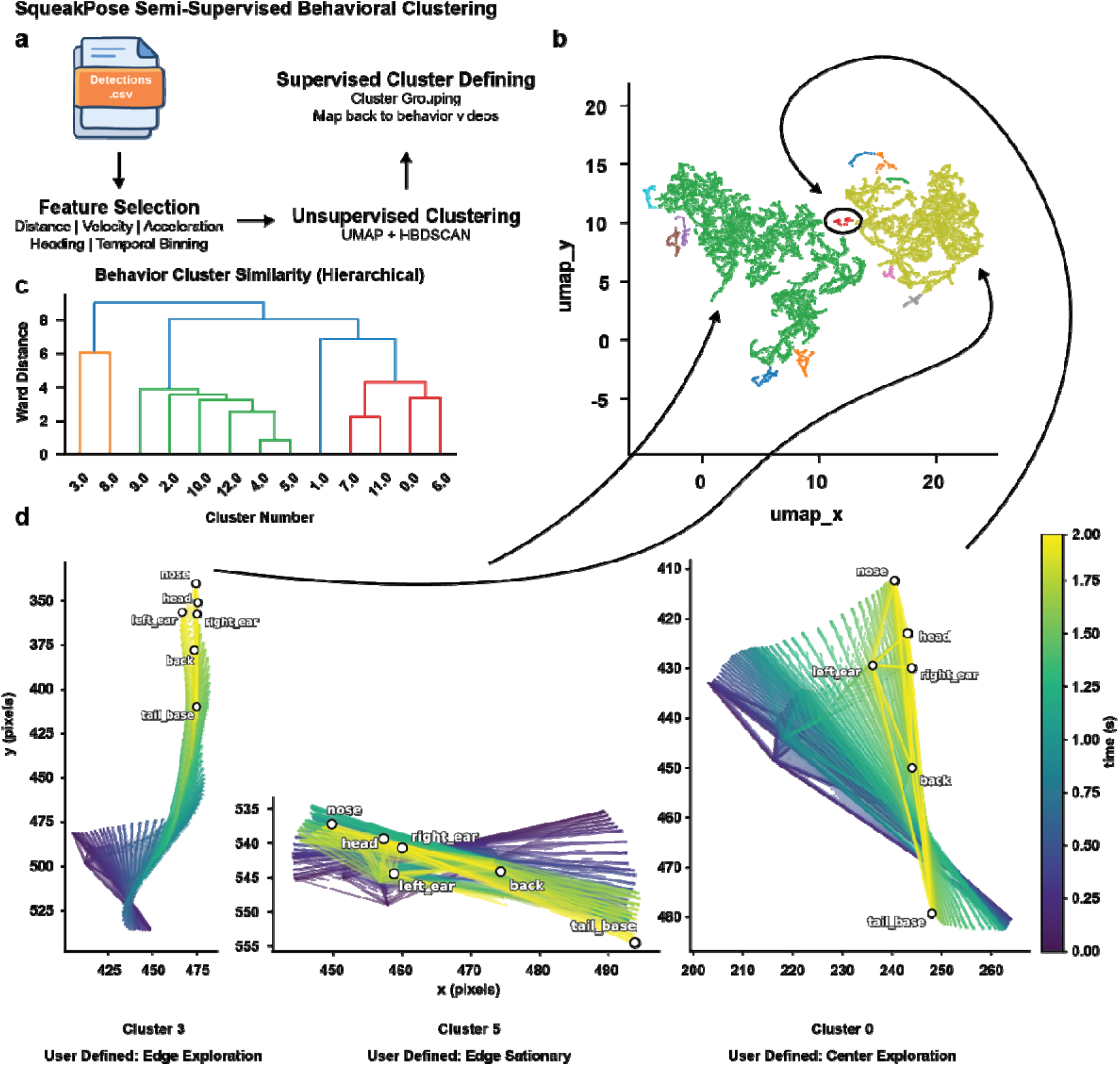
A semi-supervised workflow for identifying recurring behavioral clusters from pose-derived motion features. **(a)** Overview of the analysis pipeline. Distance, velocity, acceleration, heading, and temporal features are extracted from detection files and embedded using UMAP, followed by unsupervised clustering wit HDBSCAN. **(b)** Two-dimensional UMAP embedding of motion features colored by cluster identity, illustrating separable groups of behavioral states. **(c)** Hierarchical dendrogram showing similarity relationships among clusters using Ward distance. **(d)** Representative pose-estimation track trajectories (2 seconds total) from selected clusters, user-defined as “edge exploration,” “edge stationary,” and “center exploration.” This semi-supervised workflow provides a practical example of behavior-space discovery and labeling, with cluster outputs directly exportable to advanced models such as CEBRA or keypoint-MoSeq for further analysis.

To complement the SqueakPose Studio software ecosystem, we designed MouseHouse, a compact, hardware platform for synchronized video acquisition and behavioral control (Fig. 5a–i). While established frameworks such as DeepLabCut and SLEAP can interface with behavioral hardware through auxiliary pipelines, these implementations often rely on external middleware such as Bonsai, which provides powerful real-time control but lacks mature support for embedded Linux/ARM platforms like the Nvidia Jetson series. We built MouseHouse to achieve tighter integration between the software environment (SqueakView) and embedded hardware (MouseHouse). MouseHouse and SqueakView were co-developed to share a unified data format, timing architecture, and control interface, enabling deterministic communication between model inference, sensor inputs, and task logic.

**Figure 5 -.**
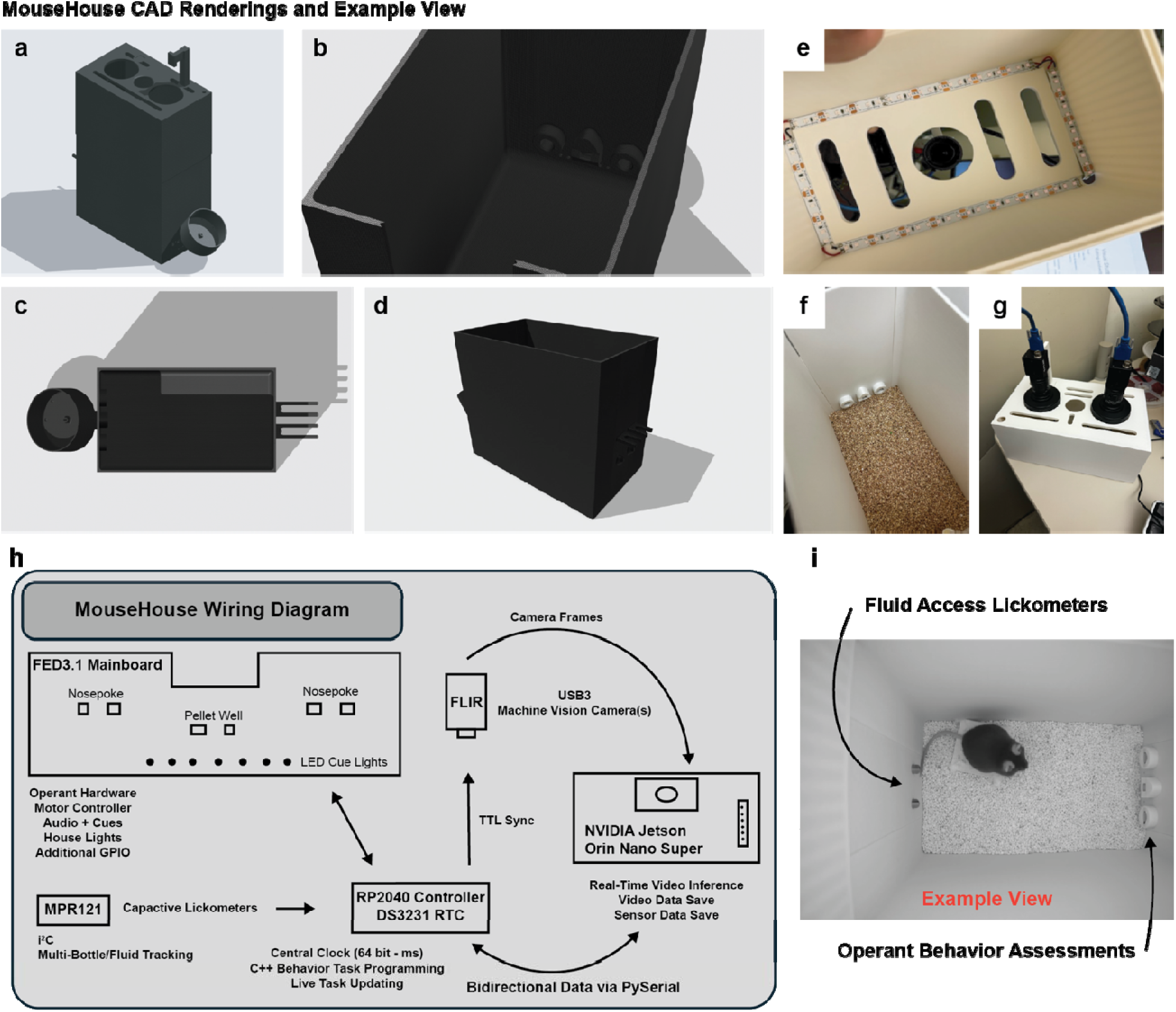
MouseHouse: A 3D-printed, edge-computing environment integrating synchronized video acquisition and behavioral hardware control. **(a–d)** Computer-aided design (CAD) renderings showing the modular enclosure, camera mount, and removable front panel. **(e)** Top-mounted USB3 machine-vision camera with visible and 850 nm infrared (IR) LED illumination for light and dark recordings. **(f–g)** Configurable arena fixtures for pellet wells and capacitive lickometers supporting fluid-access or operant tasks. **(h)** System wiring diagram linking the NVIDIA Jetson Orin Nano Super (6-core ARM CPU, 1024 CUDA cores, 32 tensor cores, 8 GB LPDDR5, 67 INT TOPS) to an RP2040 controller and MPR121 capacitive sensors. **(i)** Example configuration showing a two-bottle choice and fixed-ratio 1 (FR1) feeding task. Each unit operates as an independent edge device performing on-board YOLO inference, sensor synchronization via TTL, and bidirectional task control—enabling scalable, low-cost, networked behavioral experiments without workstation-grade GPUs.

The MouseHouse enclosure was fabricated from 3D-printed components designed in CAD (Fig.5a–d) and optimized for top-down video acquisition. The structure includes integrated mounts for USB3 machine-vision cameras that support hardware TTL triggering for frame-accurate synchronization. Uniform illumination and internal cable routing minimize visual obstruction (Fig.5e). The lighting system incorporates both visible and 850 nm infrared LEDs, enabling continuous video acquisition under standard lighting or in complete darkness. Although the standard configuration uses a single top-down camera, the design supports multi-camera stereo imaging for depth-based reconstruction or multi-angle tracking. The interior can be configured with removable bedding or task-specific inserts for fluid or pellet access (Fig.5f,g).

At the core of the system is an NVIDIA Jetson Orin Nano Super developer kit paired with an RP2040 microcontroller (Fig.5h). The Orin Nano Super features a 6-core Arm Cortex-A78AE CPU, 1024 Ampere-architecture CUDA cores, and 32 Tensor Cores, with 8 GB LPDDR5 memory (102 GB/s), delivering up to 67 INT8 TOPS of AI performance. This compute capacity enables real-time YOLO inference and full-resolution video capture through TensorRT acceleration. The RP2040 manages peripheral I/O – including capacitive-based lickometers^30^, TTL synchronization, and auxiliary hardware such as FED3^31^ controllers or cue lights and audio and communicates with the Jetson over USB serial for bidirectional data exchange and sub-millisecond synchronization between behavioral events and video frames. By using an edge-computing architecture, each MouseHouse unit contains its own GPU-enabled computer, removing the need for a centralized workstation or high-end desktop GPUs. This configuration enhances scalability, allowing multiple boxes to operate independently while being networked to a central data repository for remote monitoring, synchronization, and storage.

An example configuration is shown in Fig. 5i, where a mouse performs a two-bottle choice task using capacitive-based lickometers for fluid access, while an operant panel delivers food pellets on a fixed-ratio 1 (FR1) schedule to measure feeding behavior. The same platform can be adapted for other operant paradigms, free exploration, or multimodal sensor recording. In contrast to commercial systems that require proprietary hardware and can be prohibitively expensive, MouseHouse can be assembled from off-the-shelf components at a fraction of the cost, lowering the barrier to implementation and replication. MouseHouse delivers a modular, scalable environment for real-time, closed-loop behavioral experiments with tightly coupled software and hardware control.

To enable real-time behavioral experiments on embedded systems, we developed SqueakView, a software suite that runs natively on the Jetson Orin Nano Super and provides complete control over model deployment, video acquisition, and sensor integration (Fig.6a–f). Models trained in SqueakPose Studio can be exported as quantized TensorRT engines directly on the device through an integrated Data Engine Builder utility, which converts Ultralytics YOLO weights to ONNX and generates optimized inference engines in FP32, FP16, or INT8 precision formats (Fig.6a).

A configurable experiment-start GUI manages camera initialization, file paths, and session parameters, including resolution, frame rate, pixel format, and compression settings, while also handling serial communication with the RP2040 microcontroller (Fig.6b). During acquisition, SqueakView displays a live video stream with real-time model predictions – bounding boxes and keypoints – alongside performance indicators for CPU, GPU, RAM, and disk utilization, and synchronized sensor dashboards that visualize licks, pokes, and pellet deliveries in real-time (Fig.6c–e). For additional details and a live video demonstration of all SqueakView features, please refer to the project’s GitHub repository.

At the end of each session, all files are automatically organized in a unified directory structure compatible with the SqueakPose Analysis Utility used in Figs.3–4. Each run produces a raw video, an annotated inference video, a JSON metadata file describing model performance and camera settings, and CSV outputs for detections and sensor events (Fig.6f). This ensures full compatibility across training, inference, and analysis stages.

Performance benchmarks on the Jetson Orin Nano Super demonstrate that both detection and pose models achieve real-time inference rates sufficient for closed-loop operation, with frame latencies of approximately 7–10 ms (INT8), 15–20 ms (FP16), and 20–25 ms (FP32) (Fig. 6g–h). These results confirm that on-device inference is fast enough to support real-time behavioral labeling and task feedback without requiring extensive frame buffering or external workstation grade GPUs.

**Figure 6 -.**
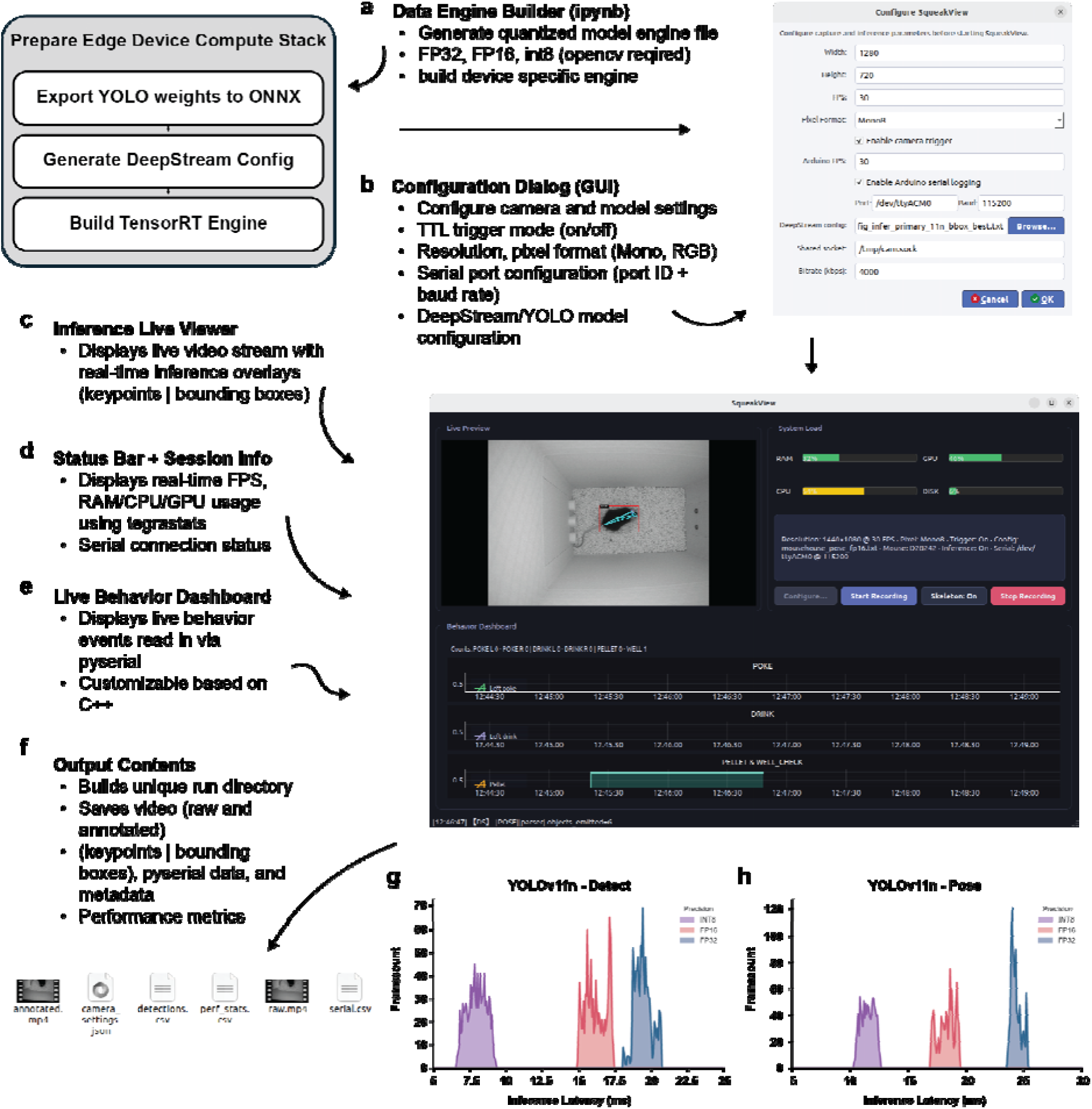
SqueakView: A graphical interface for live behavioral recording, model deployment, and performance monitoring on Jetson edge devices. **(a)** The Data Engine Builder notebook generates optimize TensorRT engines (FP32, FP16, or INT8) from trained YOLO weights for device-specific deployment. **(b)** Configuration dialog for experiment setup, including camera resolution, frame rate, pixel format, TTL trigger mode, serial communication port, and DeepStream/YOLO model selection. **(c–e)** Live interface showing real-time video fee with overlaid keypoints and bounding boxes, resource utilization (CPU, GPU, RAM), and behavior-event dashboards synchronized via serial input. (f) Output files saved per session include raw and annotated videos, JSON session metadata, and CSV logs for detections and system performance. **(g–h)** Frame-latency benchmarks for YOLOv11n detection and pose models showing real-time inference performance on the Jetson Orin Nano Super without frame buffering. Together, SqueakView enables fully embedded, real-time video analysis synchronized with behavioral sensor data.

## Discussion

Quantitative behavioral analysis has advanced rapidly over the past decade, driven by deep learning–based approaches that enable markerless tracking of animal pose and movement. Yet despite the success of frameworks such as DeepLabCut, SLEAP, and LightningPose^15^, the widespread adoption of these tools has been limited by the technical separation between model training, inference, and experimental control. Most existing systems require workstation-grade GPUs, custom data wrangling, or external middleware to link behavior with hardware for real-time deployment. As behavioral neuroscience moves toward increasingly multimodal and high-throughput paradigms, a major challenge lies in developing integrated platforms that combine modern computer vision, real-time control, and accessible hardware infrastructure.

SqueakPose Studio, MouseHouse, and SqueakView were designed to address this gap by unifying behavioral tracking, analysis, and hardware integration within a single, modular ecosystem. SqueakPose Studio builds upon recent advances in transformer-augmented YOLO architectures, providing a fast and flexible foundation for pose estimation that eliminates the need for patch-based sampling and slow post-processing pipelines. The inclusion of SqueakPose Studio and SqueakPose Analysis Utility enables users to move seamlessly from labeling to training, inference, and behavioral quantification within one standardized workflow. On the hardware side, MouseHouse extends these capabilities to the experimental arena, embedding high-speed inference and synchronized sensor logging directly within the behavioral apparatus. This pairing creates a reproducible end-to-end environment – from video acquisition to semi-supervised clustering – that can operate entirely on affordable, embedded hardware.

The decision to implement the platform on the Jetson Orin Nano Super reflects a broader architectural shift toward edge computing in neuroscience. By distributing inference to individual devices, MouseHouse eliminates the dependency on centralized workstations, lowers overall cost, and enables laboratories to scale behavioral throughput with minimal infrastructure. Each unit operates autonomously and can be networked to a shared data repository, allowing for remote monitoring and coordination of experiments. This design philosophy mirrors trends in autonomous systems and robotics, where on-device intelligence and modular sensing architectures have replaced centralized processing. Importantly, real-time inference on the Jetson platform is fast enough to support millisecond-resolution feedback, opening the door to closed-loop paradigms in which behavioral detections can trigger dynamic task updates, optogenetic stimulation, or reward contingencies.

Rather than competing with existing pose-estimation frameworks, SqueakPose Studio complements them by emphasizing deployment, usability, and integration. Tools such as DeepLabCut and SLEAP excel at training high-accuracy models for offline analysis, while SqueakPose Studio focuses on accelerating the full experimental cycle—from labeling to real-time deployment. This approach rethinks not only how behavioral data are collected, but also how they can be used dynamically during experiments. The modular design and open python interfaces further ensure compatibility with established neuroscience tools, enabling downstream synchronization with *in vivo* electrophysiology, fiber photometry, or other data streams.

Despite the progress in pose estimation, behavioral neuroscience still lacks robust frameworks for interpreting the high-dimensional datasets these models produce. Early explorations using vision-language models and embedding techniques point toward richer representations of behavior, yet these approaches remain primarily descriptive. What is ultimately needed are multimodal, closed-loop models that integrate video, sensor, and contextual data—akin to the sensor-fusion architectures developed for self-driving systems—to infer behavioral state, predict intent, and guide experimental actions in real time^32^. Achieving this level of integration will require complete control over data collection, formatting, and timing. By embedding inference, sensing, and control within a single hardware–software stack, SqueakPose Studio, MouseHouse, and SqueakView establish the foundational infrastructure for developing and testing such multimodal reasoning models in behavioral neuroscience.

Although developed and validated primarily in mice, the SqueakPose framework is model-agnostic and supports pretrained YOLO keypoint networks such as COCO human pose models^33^. When deployed on the same platform, these models can perform zero-shot tracking of human posture and movement, demonstrating the flexibility of the underlying architecture. This cross-domain compatibility positions the system as a bridge between preclinical and clinical behavioral analysis – enabling comparable, high-resolution movement quantification across species. Such translational continuity may facilitate future applications in healthcare research, where automated, real-time behavioral metrics are increasingly valuable for assessing neurological or psychiatric function.

Together, these platforms represent a shift toward accessible, edge-based experimentation, where real-time inference, multimodal integration, and closed-loop feedback become standard components of behavioral research. As AI methods evolve from static analysis to adaptive, reasoning-based systems, the ability to deploy them directly within behavioral environments will be critical for uncovering how animals perceive, decide, and act in complex contexts.

## Methods

### Animals

All animal experiments were performed with 8- to 10-week-old male and female C57BL/6J mice (Jackson Laboratory) housed in the National Institute on Alcohol Abuse and Alcoholism (NIAAA) animal facility under a standard 12 h light/dark cycle. Animals had *ad libitum* access to food and water in their homecage environments. Sex determination was limited to an assessment of external genitalia. All procedures were approved by the NIAAA Animal Care and Use Committee (protocol LIN-DL-1) and complied with the National Institutes of Health Guide for the Care and Use of Laboratory Animals.

### Software Development, Model Training, and Benchmarking

All SqueakPose software modules were written in Python 3.12 and executed within a reproducible environment defined using uv.toml configuration files (following license approval, environment files will be deposited in public repositories). Development and model training were conducted on an Apple M2 Pro Mac Mini (macOS Sequoia 15.7, 32 GB unified memory) and a custom-built Windows 11 workstation equipped with an Intel i7-8700K CPU, 32 GB RAM, and an NVIDIA RTX 4060 Ti GPU (16 GB VRAM). Training via SqueakPose Studio used PyTorch 2.9.0 (MPS-accelerated on macOS or CUDA 13.0 on Windows), Ultralytics YOLO v8.3.218, and PyQt6 6.10.0.

YOLOv11 models were trained for 150 epochs with a batch size of 8 using the Ultralytics Python libraries. Standard augmentations included random rotation (±15°), horizontal flip, scaling (0.8–1.2), and brightness/contrast jitter. A common dataset consisting of 15 videos (10 min open-field tasks of single mice with bilateral photometry cables) was used for training, validation, and testing. A total of 150 manually labeled frames (bounding boxes and keypoints) were generated in SqueakPose Studio and split 80/20 (train/validation). The same dataset was converted to DeepLabCut format to benchmark YOLO-based SqueakPose models against ResNet-50 backbones trained in DeepLabCut (Extended Data Fig. 1). For benchmarking inference times, the same full 10-minute video was predicted on all frames.

### SqueakPose Analysis Utility

Pose-derived motion features including distance, velocity, acceleration, and heading orientation were computed using the SqueakPose Analysis Utility, implemented in a Jupyter Notebook environment. Coordinates were converted to real-world units via a built-in calibration GUI using video metadata extracted with OpenCV. A 1-Euro smoothing filter was applied to raw tracking coordinates to suppress high-frequency noise.

Feature embeddings were generated using UMAP v0.5.9 (n_neighbors = 50, min_dist = 0.3, spread = 1, metric = ‘euclidean’, random_state = 42) and clustered with HDBSCAN v0.8.40 (min_cluster_size = 40, min_samples = 10, metric = ‘euclidean’, cluster_selection_epsilon = 0.3, cluster_selection_method = ‘leaf’). Hierarchical relationships among clusters were computed using Ward linkage. Resulting feature matrices and cluster assignments were visualized in Matplotlib and Seaborn and exported as .csv files for downstream use. Processed dataframes can be converted to NWB, parquet, or other compressed formats via standard I/O nodes for long-term archival or integration with tools such as CEBRA or Keypoint-MoSeq.

### MouseHouse Hardware

Each MouseHouse unit consisted of a 3D-printed behavioral arena with integrated electronics for real-time video acquisition and control. Designs were created in Shapr3D, exported as STEP files, and printed in matte white PLA (Bambu Labs) on a Bambu Lab X1 Carbon printer (0.2 mm layer height, 20 % infill). The assembled enclosure measured approximately 10 × 6 × 14 inches (L × W × H) and included custom mounts for the camera, IR illumination, operant hardware, lickometer-enabled drinking tubes, and controller board. CAD and STEP files for all printed parts are publicly available at https://github.com/dlhagger/squeakview.

### Video Acquisition and Optical Assembly

Video acquisition used a FLIR Blackfly S BFS-U3-16S2M-CS monochrome CMOS camera (USB 3.1, 1.6 MP). Cameras were fitted with 4 mm UC-Series C-mount lenses (Edmund Optics #33300) and UV/VIS-cut filters (Edmund Optics #89842) to restrict imaging to near-infrared wavelengths. Cameras connected to the Jetson Orin Nano Super via 3 m locking USB 3.1 cables and received TTL triggers from an Adafruit Feather RP2040 Adalogger microcontroller for frame-accurate synchronization. Uniform illumination was provided by 850 nm IR LED strips (Waveform Lighting, USA) mounted along the inner cage lid and powered via the 5 V header.

### Embedded Control and Synchronization

Behavioral control, event logging, and timing synchronization were managed by an Adafruit Feather RP2040 Adalogger microcontroller running custom firmware written in Arduino C. The RP2040 served as the master clock for all timing and maintained precision via a DS3231 Precision RTC FeatherWing (±2 ppm). The Feather stack interfaced directly with the FED3 breakout board (Open Ephys OEPS-7511). Firmware modifications extended the FED3 library with a motor-controller state machine and non-blocking timing logic to minimize latency and drift. Capacitive sensing for fluid intake was handled via Adafruit MPR121 12-key capacitive touch sensors, communicating with the RP2040 over I²C.

All events were timestamped using a 64-bit microsecond counter on the RP2040. Timestamp data were transmitted via serial using pySerial v3.5 to the Jetson Orin Nano Super, which appended corresponding Unix timestamps for clock alignment. TTL triggers on dedicated GPIO pins synchronized behavioral events with video frame acquisition. The system architecture designates the RP2040 as the timing master because its clock precision and GPIO latency outperform the Jetson’s system timer.

### Edge Inference and MouseHouse Data Acquisition

SqueakView provides the operator interface for video acquisition, model deployment, and behavioral event logging within the MouseHouse environment. It runs on a Jetson Orin Nano Super (Ubuntu 22.04, JetPack 6.2.1, CUDA 12.6) and coordinates three isolated uv-managed environments for capture, DeepStream inference, and GUI control.

The capture module configures the FLIR Blackfly camera (resolution, FPS, exposure, trigger mode) using PySpin and streams frames to shared memory via a custom GStreamer pipeline. The inference module consumes this stream and executes inference using TensorRT engines; performance metrics are recorded through the NvDsPerformanceStruct API in the NVIDIA DeepStream SDK v7.1. The GUI provides a configuration dialog (capture parameters, DeepStream config, serial settings), displays a live preview with Jetson resource usage, and visualizes TTL-derived behavioral counters. A backend controller manages process launch, logging, and automatic organization of session outputs, including raw video, annotated frames, detection .csv files, and JSON metadata.

### TensorRT Deployment and Engine-Build Workflow

YOLO deployment followed a unified TensorRT build framework implemented within the DeepStream-YOLO repository, which organizes model artifacts into a structured hierarchy (weights, ONNX exports, DeepStream configs, and serialized engines). Model conversion proceeded through a notebook-driven pipeline that abstracts the PyTorch → ONNX → TensorRT toolchain into a deterministic workflow built on device for correct paramterization.

Within this framework, YOLO weights were exported to ONNX, automatically integrated into a templated DeepStream configuration, and compiled into optimized FP32, FP16, or INT8 TensorRT engines. INT8 builds used calibration imagery from in-house mousehouse datasets to correct for errors in the INT8 quantization. The pipeline additionally produced all auxiliary files required for deployment, including DeepStream configuration files, parser settings, and device-specific metadata, matched to the target Jetson architecture to ensure compatibility and stable runtime execution. Multiple precision builds can coexist within standardized directories, enabling consistent deployment by the SqueakView inference subsystem.

### Data, Code and Materials Availability

All source code, CAD files, firmware, hardware build files and software environment specifications generated in this study are publicly accessible at https://github.com/dlhagger/SqueakPoseStudio and https://github.com/dlhagger/squeakview. The datasets supporting the findings of this work are available from the corresponding author (David L. Haggerty) upon reasonable request.

As allowed by the licenses associated with the SqueakPose and MouseHouse GitHub repositories, the software and associated materials are freely available for academic and non-profit research use, while commercial use may require a separate software license from the NIH. All use, distribution, duplication, and modification must additionally comply with the licensing terms of third-party software dependencies incorporated into the system.

A live video demonstration illustrating the full workflow and principal features of the system is provided in the README files of the respective GitHub repositories.

## Acknowledgements

“This research was supported by the Intramural Research Program of the National Institutes of Health (NIH). The contributions of the NIH author(s) are considered Works of the United States Government. The findings and conclusions presented in this paper are those of the author(s) and do not necessarily reflect the views of the NIH or the U.S. Department of Health and Human Services.”

## Acknowledgements

This work was supported by the intramural program of NIAAA, NIH 1ZIAAA000407-22 (DML) and the PRAT fellowship program of NIGMS, NIH 1FI2GM154674-01 (DLH).

## References

1. Saad Saoud, L., et al. Beyond observation: Deep learning for animal behavior and ecological conservation. Ecological Informatics 84, 102893 (2024).

2. Wei, K. & Kording, K. P. Behavioral tracking gets real. Nat Neurosci 21, 1146–1147 (2018).

3. Mathis, M. W. & Mathis, A. Deep learning tools for the measurement of animal behavior in neuroscience. Current Opinion in Neurobiology 60, 1–11 (2020).

4. Perez, M. & Toler-Franklin, C. CNN-Based Action Recognition and Pose Estimation for Classifying Animal Behavior from Videos: A Survey. Preprint at 10.48550/arXiv.2301.06187 (2023).

5. Grieco, F. et al. Measuring Behavior in the Home Cage: Study Design, Applications, Challenges, and Perspectives. Front. Behav. Neurosci. 15, (2021).

6. Couzin, I. D. & Heins, C. Emerging technologies for behavioral research in changing environments. Trends Ecol Evol 38, 346–354 (2023).

7. Borowiec, M. L. et al. Deep learning as a tool for ecology and evolution. Methods in Ecology and Evolution 13, 1640–1660 (2022).

8. Bertram, M. G. et al. Frontiers in quantifying wildlife behavioural responses to chemical pollution. Biological Reviews 97, 1346–1364 (2022).

9. Luxem, K. et al. Open-source tools for behavioral video analysis: Setup, methods, and best practices. eLife 12, e79305 (2023).

10. Kennedy, A. The what, how, and why of naturalistic behavior. Current Opinion in Neurobiology 74, 102549 (2022).

11. Marshall, J. D., Li, T., Wu, J. H. & Dunn, T. W. Leaving flatland: Advances in 3D behavioral measurement. Current Opinion in Neurobiology 73, 102522 (2022).

12. Pereira, T. D., Shaevitz, J. W. & Murthy, M. Quantifying behavior to understand the brain. Nat Neurosci 23, 1537–1549 (2020).

13. Mathis, A. et al. DeepLabCut: markerless pose estimation of user-defined body parts with deep learning. Nature Neuroscience 21, 1281–1289 (2018).

14. Pereira, T. D. et al. SLEAP: A deep learning system for multi-animal pose tracking. Nature Methods 19, 486–495 (2022).

15. Biderman, D. et al. Lightning Pose: improved animal pose estimation via semi-supervised learning, Bayesian ensembling and cloud-native open-source tools. Nat Methods 21, 1316–1328 (2024).

16. He, K., Zhang, X., Ren, S. & Sun, J. Deep Residual Learning for Image Recognition. Preprint at 10.48550/arXiv.1512.03385 (2015).

17. Bonsai. *GitHub* https://github.com/bonsai-rx.

18. Kane, G. A., Lopes, G., Saunders, J. L., Mathis, A. & Mathis, M. W. Real-time, low-latency closed-loop feedback using markerless posture tracking. eLife 9, e61909 (2020).

19. Khanam, R. & Hussain, M. YOLOv11: An Overview of the Key Architectural Enhancements. Preprint at 10.48550/arXiv.2410.17725 (2024).

20. Tian, Y., Ye, Q. & Doermann, D. YOLOv12: Attention-Centric Real-Time Object Detectors. Preprint at 10.48550/arXiv.2502.12524 (2025).

21. Li, G., Jian, R., Jun, X. & Shi, G. A Review of You Only Look Once Algorithms in Animal Phenotyping Applications. Animals (Basel) 15, 1126 (2025).

22. Niko Sirmpilatze et al. neuroinformatics-unit/movement: v0.10.0. Zenodo 10.5281/ZENODO.12755724 (2025).

23. Frontiers | PainSeeker: a head pose-invariant deep learning method for assessing rat’s pain by facial expressions. https://www.frontiersin.org/journals/veterinary-science/articles/10.3389/fvets.2025.1619794/full.

24. DeepLabCut/DLCutils. DeepLabCut (2025).

25. DeNardoLab/BehaviorDEPOT. DeNardoLab (2025).

26. mlfpm/deepof. MLFPM (2025).

27. Seibenhener, M. L. & Wooten, M. C. Use of the Open Field Maze to Measure Locomotor and Anxiety-like Behavior in Mice. J Vis Exp 52434 (2015) doi:10.3791/52434.

28. Schneider, S., Lee, J. H. & Mathis, M. W. Learnable latent embeddings for joint behavioural and neural analysis. Nature 617, 360–368 (2023).

29. Weinreb, C. et al. Keypoint-MoSeq: parsing behavior by linking point tracking to pose dynamics. Nat Methods 21, 1329–1339 (2024).

30. Peripheral alcohol metabolism dictates ethanol consumption and drinking microstructure in mice - Mackowiak - 2025 - Alcohol, Clinical and Experimental Research - Wiley Online Library. https://onlinelibrary.wiley.com/doi/10.1111/acer.70036.

31. Matikainen-Ankney, B. A. et al. An open-source device for measuring food intake and operant behavior in rodent home-cages. eLife 10, e66173 (2021).

32. Alpamayo-R1: Bridging Reasoning and Action Prediction for Generalizable Autonomous Driving in the Long Tail | Research. https://research.nvidia.com/publication/2025-10_alpamayo-r1.

33. Ultralytics. COCO. https://docs.ultralytics.com/datasets/pose/coco.

