## Supplementary material for "An end-to-end platform for pose estimation and real-time edge-AI deployment": figure 1 - figure 1 supplement figure legend text

**Figure 1-figure supplement 1 -** *Training and validation performance of YOLOv11s-pose and DeepLabCut (ResNet-50) models.* (a–b) Normalized training and validation loss across 150 epochs for YOLOv11s-pose (orange) and DeepLabCut (DLC; gray) trained on identical datasets of 150 manually labeled frames from open-field recordings. Both models achieved stable convergence with low final loss values. (c–d) Mean Average Precision (mAP) at IoU = 0.5 (mAP50) and averaged across IoU = 0.5–0.95 (mAP50-95) plotted across training epochs. YOLOv11s-pose achieved rapid convergence and maintained mAP values comparable to DeepLabCut throughout training, demonstrating equivalent pose-estimation accuracy under matched conditions.
